# Deep shifts in Evolutionary Rate Trajectories of Ancient Bacterial Genes

**DOI:** 10.64898/2026.09.09.750377

**Authors:** Hayley B. Hassler, Tanisha Gupta, Anton S. Petrov, Loren Dean Williams, Claudia Alvarez-Carreño

## Abstract

Reconstruction of ancestral gene repertoires from extant genomes captures only the genes that survived; those lost from the record are invisible. Among the genes that did persist, selective pressures change not only across lineages but across time.

Here, we resolve evolutionary rate trajectories across 528 genes in the Last Bacterial Common Ancestor (LBCA). We hypothesized that LBCA genes would show distinct evolutionary rate trajectories across bacterial history and tested this by resolving normalized branch lengths across five calibrated taxonomic intervals (phylum, class, order, family, and genus), spanning approximately 2.1 billion years of bacterial diversification.

The distribution of rate trajectories is continuous, but four clusters capture the major patterns: Decelerating, Class-Peaking, Constant, and Accelerating. The Decelerating and Class-Peaking clusters are composed predominantly of Genetic Information Processing genes, whereas the Constant and Accelerating clusters are enriched for Metabolic genes. Specifically, the Decelerating cluster is enriched for core informational machinery, including translation initiation factors and components of the expressome, the molecular complex physically coupling transcription and translation, suggesting that transcription-translation interfaces locked in early in bacterial history. Cofactor-dependence and cofactor biosynthesis are decoupled: biosynthetic pathways producing metallocofactors such as heme, molybdopterin, and cobalamin, concentrated in the Constant and Accelerating clusters but proportionally more proteins use inorganic cofactors in the Decelerating and Class-Peaking clusters. This offset coincides with the shift in metal bioavailability associated with the Great Oxidation Event and reveals genomic fingerprints of the co-evolution of bacterial metabolism with planetary geochemistry.

## Introduction

Bacteria have an evolutionary history spanning approximately 3.5 billion years and encompass the greatest genetic diversity of the three domains of life (Pace 1997; Whitman, et al. 1998; Knoll and Nowak 2017). The Last Bacterial Common Ancestor (LBCA) represents the hypothetical ancestral node of the bacterial phylogenetic tree from which all surviving bacterial lineages diverged. Genes conserved from the LBCA to the present retain signatures of deep evolutionary history, decipherable through sequence variation and substitution patterns.

To reconstruct ancestral gene repertoires and infer presence in the LBCA, we identify genes present across diverse lineages and find signals of vertical inheritance (Delaye, et al. 2005; Berkemer and McGlynn 2020; Hyun and Palsson 2023). The genes in the LBCA repertoire are, by definition, those that persisted across 3.5 billion years of divergence. Their retention in bacterial genomes reflects both historical contingency and selective pressures. Reconstruction from extant genomes captures only the genes that survived; those lost from the record are invisible.

Selective pressures to retain a gene vary across lineages at any given point in time. The GroEL chaperonin, for example, is widely distributed across bacteria (Lund 2009) and essential in *Escherichia coli* yet absent from some species of mollicutes (Clark and Tillier 2010). Similarly, essentiality and retention of ribosomal protein genes appear to be lineage-specific (Galperin, et al. 2021; Nikolaeva, et al. 2021; Mise and Iwasaki 2022).

The magnitudes of selective forces can be inferred from rates of sequence change (Wilson, et al. 1977). Genes under negative selection evolve slowly and accumulate fewer substitutions. Consistently, studies have found that essential genes evolve more slowly than nonessential genes (Jordan, et al. 2002; Rocha and Danchin 2004; Zhang and He 2005; Luo, et al. 2015). These rate estimates are typically calculated as a single average over a fixed evolutionary depth, collapsing variation in selective pressure over time into one number. But selective pressures change not only across lineages but across time, and we hypothesized that genes would show distinct evolutionary rate trajectories across bacterial history.

Diversification of bacterial species is driven by processes ranging from the emergence of major evolutionary innovations to adaptation to differing nutritional and environmental constraints. Many of these processes are governed by lineage-restricted genes that reflect history within a certain phylogenetic group. In the current study, we use the universal bacterial gene set as a proxy for tracing evolutionary trajectories. These genes are retained across the full breadth of bacterial diversity and provide a record from the LBCA to extant lineages. By tracking divergence recorded onto the universal bacterial genes, we can reconstruct how major evolutionary events were imprinted onto the genes that carry out essential functions across bacterial cells. This approach treats persistent gene sets as a molecular archive of the selective and adaptive pressures that have acted on bacteria throughout their evolutionary history.

To evaluate changes in evolutionary rates of individual bacterial genes over time, we calculated the extent of evolutionary change across different temporal depths as defined by taxonomic level (phylum, class, order, family, and genus). We used branch lengths (measured in expected substitutions per site) as a metric for the extent of evolutionary divergence accumulated over a given time interval (Felsenstein 2004). Branch lengths capture rate variation over a gene’s history. A gene that evolves rapidly early in its history and slowly thereafter will carry a different branch length signature than one that evolves slowly early in its history and rapidly thereafter, even if both accumulate similar total change. Branch lengths are not uniform across genes spanning the same evolutionary time period; different gene sets can produce vastly different branch lengths despite representing identical timescales (Moody, et al. 2022). We therefore normalized branch lengths for comparison. Here, using normalized branch lengths we examine rate shifts that occurred deep in bacterial history, at and after the time of the LBCA.

We present a reconstruction of the LBCA gene complement, focusing on highly conserved, single-copy protein-coding genes. Our reconstruction yielded 528 genes across 2,982 genera. For each gene, we characterized evolutionary rate trajectories across five sequential taxonomic levels, and identified four gene clusters: Decelerating, Class-Peaking, Constant, and Accelerating. The Decelerating and Class-Peaking clusters are composed predominantly of Genetic Information Processing genes, whereas the Constant and Accelerating clusters are enriched for Metabolic genes. These functional differences are accompanied by a shift in cofactor usage across clusters, from a predominance of inorganic cofactors in genes in the Decelerating cluster to organic cofactors in the Accelerating cluster.

## Results

### LBCA gene content

Using 2,982 bacterial genera and 144 archaeal genera, we identified a total of 528 genes present in the LBCA (Figure 1 and Supplementary Table 1). The gene repertoire includes 302 universally distributed genes (present in 75% or more of both archaeal and bacterial genera), as well as 226 bacteria-specific genes (present in 75% or more of bacterial genera and 10% or fewer of archaeal genera).

**Figure 1.**
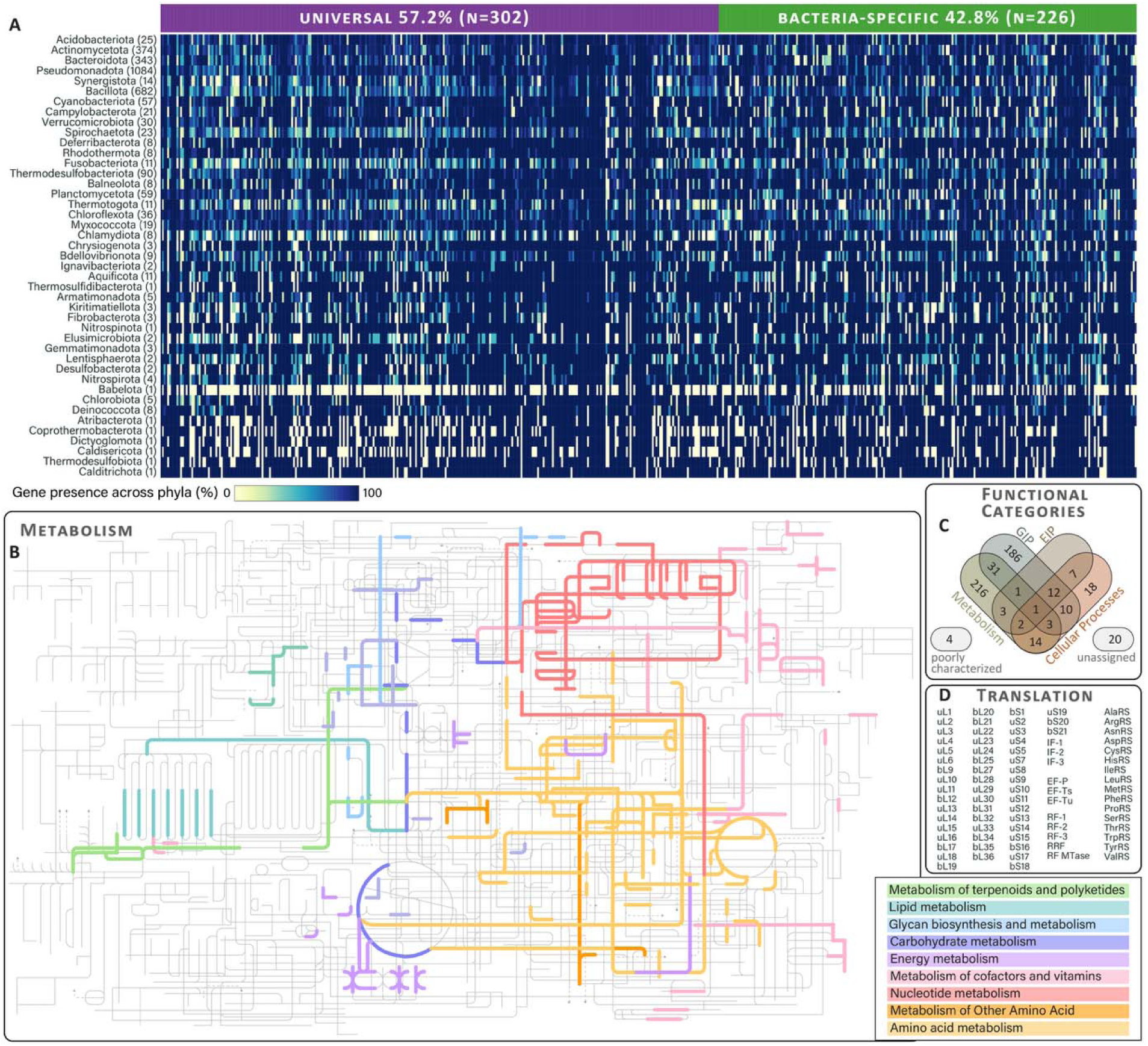
**A reconstruction of the LBCA gene** repertoire. (A) Reconstruction of the LBCA gene repertoire from gene presence across 43 bacterial phyla. Each column is one gene of 528 genes observed, and each row is a bacterial phylum. Color indicates the average proportion of genomes within that phylum carrying the gene. (B) LBCA gene repertoire mapped on to KEGG metabolic pathways. (C) Functional annotation of LBCA gene repertoire. (D) Overview of translation genes in LBCA. GIP: Genetic Information Processing. EIP: Environmental Information Processing. Metabolic pathway was generated with iPath3.0 (Darzi, et al. 2018).

### Functional characterization of LBCA genes

Genes were assigned to KEGG Orthology annotations using eggNOG-mapper v2 (Cantalapiedra, et al. 2021). A total of 495 LBCA genes (94 %) in our repertoire were assigned to KEGG (Figure 1 and Supplementary Table 1). We grouped the annotated LBCA genes by functional category, according to KEGG BRITE: Metabolism (n = 209), Genetic Information Processing (GIP, n = 184), and Cellular Processes (n = 14). 82 genes were assigned to combinations of these categories and are reported as mixed. Four genes were assigned to a KO identifier without BRITE hierarchy.

### Evolutionary rate trajectories of LBCA genes

To quantify the distribution of evolutionary change across five sequential taxonomic levels (phylum, class, order, family, and genus), we constructed an unrooted gene tree for each of the 528 LBCA genes. From each tree, we calculated the median length across all taxa at each taxonomic level. Branch lengths were estimated using only the slowest-evolving alignment sites (rate categories 1 to 5 under the LG+R10 model) to capture evolutionary signal at deep taxonomic levels and reduce saturation artifacts. Representative taxa were randomly subsampled across 100 replicates. The median across replicates was taken as the final value (see Methods). The five median values were normalized by scaling their sum to one, yielding the normalized median branch length (NMBL) at each taxonomic level for each gene.

Evolutionary rate trajectories of LBCA genes are distributed along a continuous gradient. Principal coordinate analysis (PCoA) was performed on Bray-Curtis dissimilarities among rate trajectories for all 528 genes (Figure 2B). The resulting ordination reveals a continuous U-shaped arc with no discrete gaps. PC1 (86% of variance) separates genes with peak evolutionary rates at deeper divergences (phylum, class) from those with peak evolutionary rates at recent divergences (family, genus). PC2 (10% of variance) separates genes by overall shape of the trajectory.

**Figure 2.**
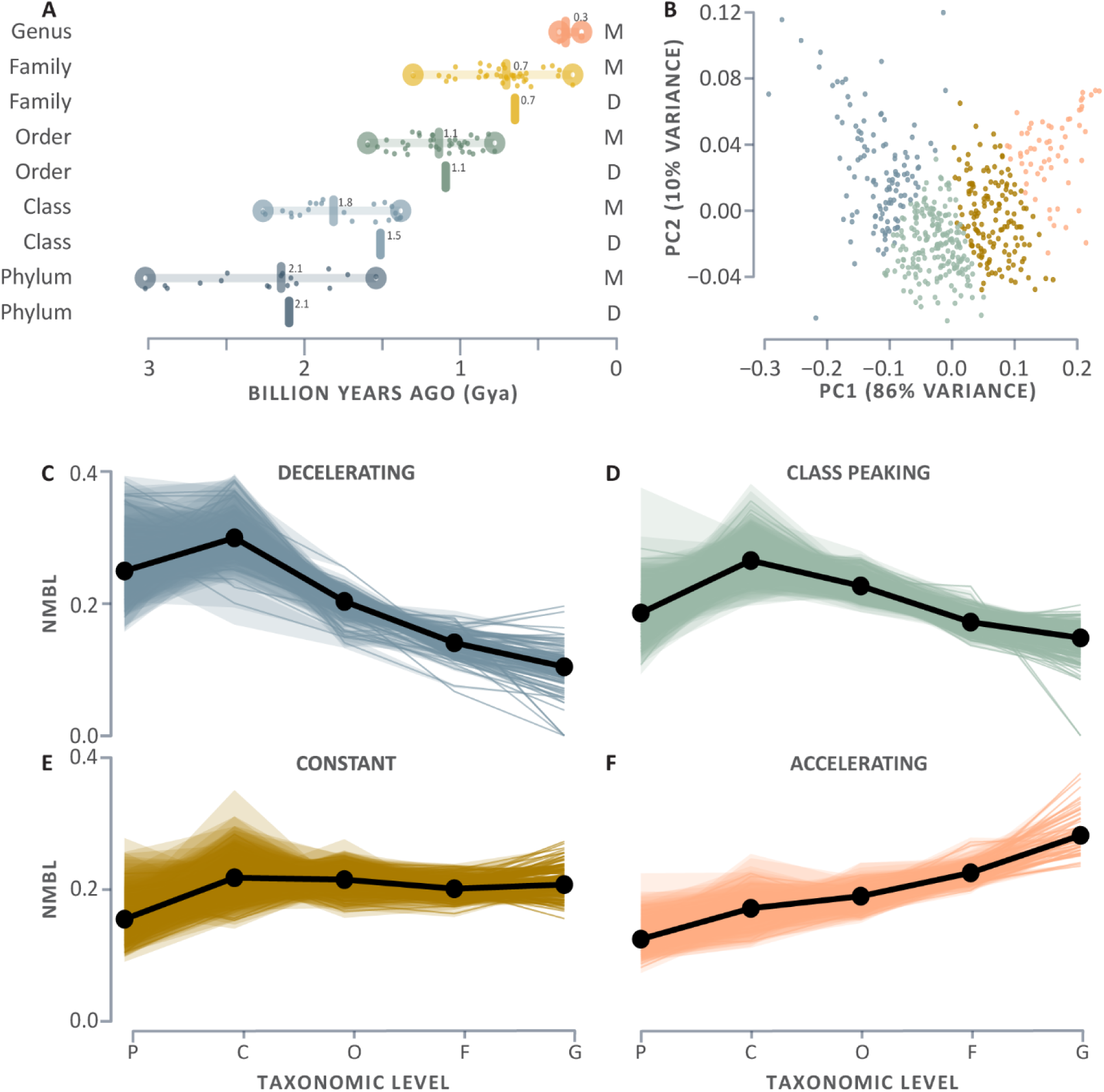
Evolutionary rate trajectories of LBCA genes. (A) Estimated divergence times for taxonomic levels from phylum to genus, based on two independent studies (M = Moody et al.; D = Davin et al.). Horizontal lines show the median divergence time for each level; individual points are divergence time estimates for individual taxonomic groups within that level. Davin et al. did not report genus-level divergence times. (B) Principal coordinate analysis of Bray-Curtis dissimilarities among evolutionary rate trajectories for all 528 LBCA genes. Each point is the median value of one gene, colored by its position along PC1. PC1 captures 86% of the variance; PC2 captures 10%. (C-F) Normalized median branch lengths across taxonomic levels for all 528 genes, organized into four clusters by k-means clustering (k=4): (C) Decelerating (n= 117); (D) Class-Peaking (n= 203); (E) Constant (n= 148), (F) Accelerating (n= 60). Black lines show the median trajectory for each cluster. Shaded ribbons show the interquartile range across 100 resampling replicates. P: phylum, C: class, O: order, F: family, G: genus. NMBL: normalized median branch length.

We applied k-means clustering to group the 528 evolutionary rate trajectories (Figure 2, Supplementary Table 1). Four clusters were identified (k=4). The clusters are: Decelerating (n= 117), Class Peaking (n= 203), Constant (n= 148), and Accelerating (n= 60) (Figure 2C-2F).

### Calibrating taxonomic levels to absolute time

We used previous estimations of the absolute time of bacterial taxonomic-level divergences (Moody, et al. 2024; Davin, et al. 2025) to calibrate our comparisons (Figure 2A): phylum-level splits at a median of 2.1 Gya (1.5 to 3.0 Gya); class-level splits at 1.5 to 1.8 Gya (0.5 to 3.0 Gya); order-level splits at 1.1 Gya (1.3 to 2.2 Gya); family-level splits at 0.7 Gya (0.3 to 1.3 Gya); and genus-level splits at a median of 0.3 Gya (0.2 to 0.4 Gya). Because the gaps between consecutive taxonomic splits in absolute time are similar in duration (∼0.45 Gya), this calibration provides a relative scale for interpreting rate trajectories. A gene evolving at a constant rate would accumulate approximately equal proportions of its total evolutionary change across each interval. With five taxonomic levels considered here, this corresponds to approximately 0.2 of the total change per split. However, taxonomic-level splits are not perfectly equidistant in absolute time, thus the 0.2 mark is used here as an estimate.

The Decelerating group shows the highest proportional rate of change at the phylum and class splits before declining steadily in rate of change through the remaining intervals (Figure 2C). The Class Peaking group follows a similar overall shape but starts from a lower rate at the phylum-level followed by a proportionally larger rate at the class-level split (Figure 2D). The Constant group shows the least overall deviation from 0.2 NMBL, with only a slightly increase at the class split (Figure 2E). The Accelerating group shows rate increase at every interval (Figure 2F).

### Functional composition of evolutionary rate clusters

The Decelerating cluster (n=117) is dominated by genetic information processing genes, including ribosomal proteins and translation factors (Figure 3A, bS1; uS2; uS3; uS4; uS5; bS6; uS7; uS8; uS9; uS10; uS11; uS12; bS18; uS19; bS21; uL1; uL2; uL4; uL5; uL10; uL11; uL14; uL16; bL17; bL19; bL20; bL28; uL29; bL31; bL32; uL33; *infA*; *infB*; *infC*; *efp*; *tuf*; *lepA*; *prfC*), transcription (Figure 3C, *rpoA*; *rpoB*; *rpoC*; *rpoZ*; *nusA*, *nusG*; *rho*; *dksA*; *greA*; *rnj*; *pnp*), tRNA biogenesis and aminoacylation enzymes (Figure 3E, ArgRS; AsnRS; AspRS; ThrRS; ValRS; IleRS; PheRS-alpha; TrpRS; MetRS (*metG*); Asp/Glu-ADT subunit B (*gatB*); *rnz*; *tgt*; *gidA*; *miaB*; *tsaD*; *dus*), and ribosome biogenesis factors (Figure 3B, *rlmH*; *rimO*; *ybeY*; *ychF*; *typA*; *hpf*; *rng*). This cluster also includes energy metabolism genes (Figure 3K), primarily subunits of the NADH:ubiquinone oxidoreductase (*nuoA*; *nuoB*; *nuoD*; *nuoI*; *nuoJ*; *nuoK*; *sdhA*; *sdhB*), succinate dehydrogenase (*sdhD*), and ATP synthase complexes (*atpA*; *atpD*; *atpE*; *atpG*).

**Figure 3.**
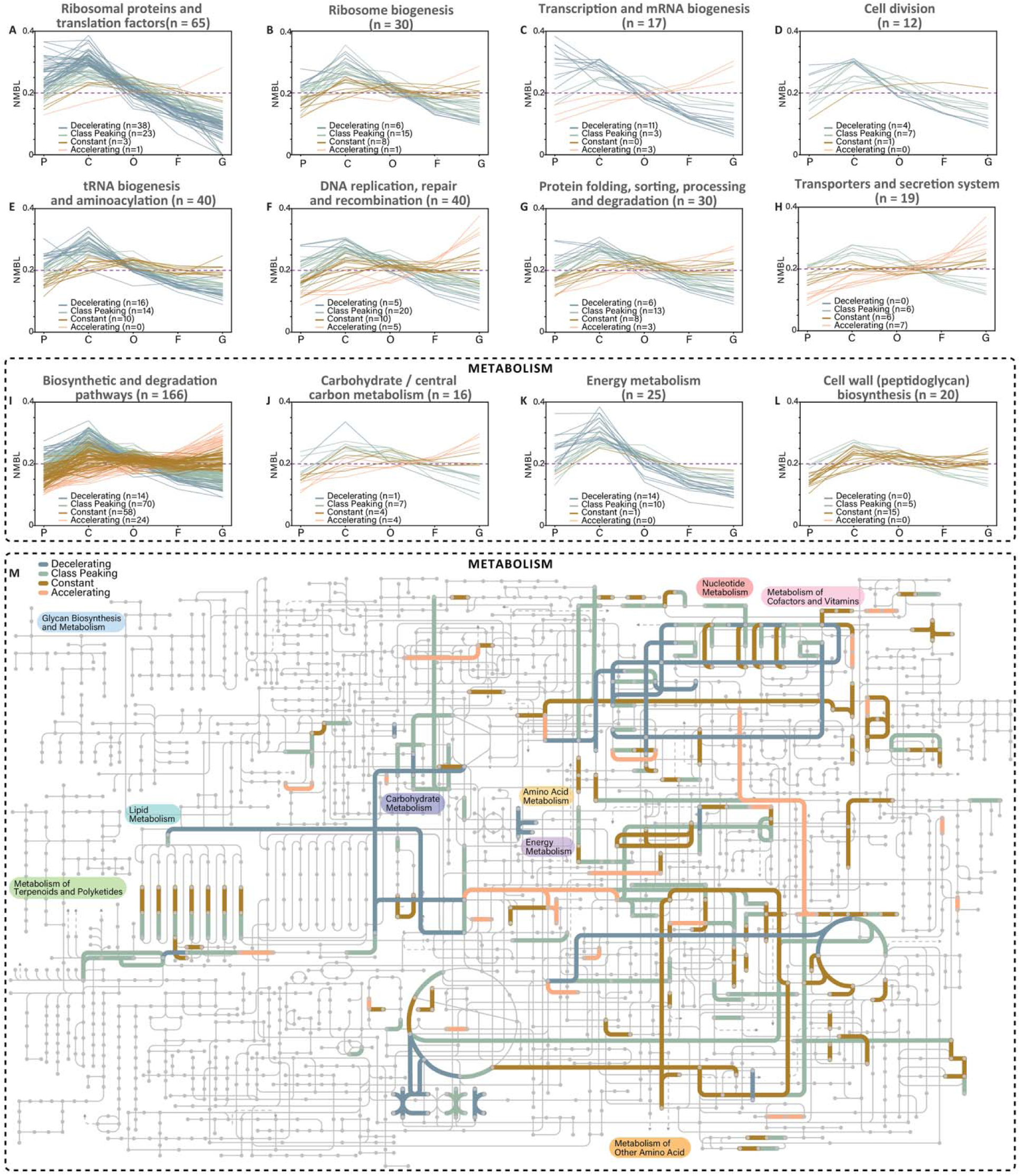
Evolutionary rate clusters within each KEGG-BRITE functional category. Normalized median branch lengths (NMBL) are shown across five sequential taxonomic levels (P: phylum, C: class, O: order, F: family, G: genus) for genes grouped by KEGG-BRITE functional category. Each line represents one gene, colored by rate-trajectory cluster. (A) Ribosomal proteins and translation factors (n=65). (B) Transporters and secretion system (n=19). (C) Transcription and mRNA biogenesis (n=17). (D) Cell division (n=12). (E) tRNA biogenesis and aminoacylation (n=40). (F) DNA replication, repair and recombination (n=40). (G) Protein folding, sorting, processing and degradation (n=30). (H) Ribosome biogenesis (n=30). (I) Biosynthetic (and degradation) pathways (n=166). (J) Carbohydrate/central carbon metabolism (n=16). (K) Energy metabolism (n=25). (L) Cell wall (peptidoglycan) biosynthesis (n=20). (M) Metabolic map showing colored by rate-trajectory clusters; grey lines indicate reactions not represented in the LBCA gene set. Metabolic pathway was generated with iPath3.0 (Darzi, et al. 2018).

The Class Peaking cluster (n=203) is the largest and most functionally diverse. It includes multiple metabolic proteins (Figure 3I), including nearly all steps of the purine (*purB*, *purD*, *purE*, *purF*, *purH*, *purL*, *purM*, *guaB*) and pyrimidine (*pyrB*, *pyrF*, *pyrH*, *carA*, *carB*) biosynthesis pathways, and nucleotide interconversion kinases (*adk*, *ndk*, *gmk*), as well as amino acid biosynthesis enzymes, including arginine (*argB*, *argC*, *argH*), branched-chain amino acids (ilvN, ilvC, leuA, leuC, lysC), histidine (*hisB*, *hisD*, *hisF*, *hisH*), tryptophan (*trpA*, *trpB*, *trpD*), and glutamate (*gltA*, *gltB*). Membrane lipid biosynthesis is also well-represented, with fatty acid synthesis enzymes (*fabD*, *fabI*, *acpP*, *accA*, *accB*, *accC*), phospholipid head-group assembly (*pssA*, *cdsA*, *pgsA*), and lipopolysaccharide acylation and lipid modification enzymes (*lpxA*, *lgt*, *lnt*, *kdsD*). It also contains DNA replication, repair, and recombination genes (Figure 3F, *polA*; *dnaE*; *holA*; *dnaX*; *gyrA*; *gyrB*; *dnaB*; *priA*; *rarA*; *topA*; *radA*; *recA*; *recF*; *recG*; *recJ*; *recN*; *recR*; *ruvA*; *ruvC*; *uvrC*), and genes related to protein folding sorting, processing and degradation (Figure 3G), including the major chaperone systems GroEL/GroES and DnaK. Cell division genes (Figure 3D, *ftsA*; *mraZ*; *ftsH*; *ftsI*; *mrp*; *scpA*; *mreC*) also concentrated here.

The Constant cluster (n=148) includes genes involved in ribosome biogenesis (Figure 3B, *rsmD*, *rsmE*, *rimP*, *rluB*, *rumA*, *rsmH*, *rsmI*, *tlyA*). Metabolic enzymes (Figure 3I) include purine salvage pathway (apt, hpt), cofactor biosynthesis pathways, including folate (*folC*, *folD*, *folP*, *metF*), coenzyme A (*coaBC*, *coaE*, *coaX*), riboflavin (*ribBA*, *ribF*, *ribH*), NAD (*nadC*, *nadD*, *nadE*), thiamine (*thiG*, thiL), pantothenate (*panB*), lipoic acid (*lipB*), acyl carrier protein activation (*acpS*), heme (*hemB*, *hemC*, *hemL*), and molybdopterin (*moaA*, *moaC*, *moaE*). Cell wall biosynthesis (Figure 3L) is also prominent, with several genes involved in peptidoglycan assembly (*ftsW*, *amiC*, *dacF*, *murE*, *murI*, *lpxD*, *mltG*, *tagO*, *uppS*, *rfbA*, *rfbB*, *ugd*, *hldE*, *ddl*, *yfiH*).

The Accelerating cluster (n=60) is the smallest. The biosynthetic component of the Accelerating cluster includes genes involved in the synthesis of folate (*folK*), biotin (*birA*), riboflavin (*ribD*), pantothenate (*panC*), FAD (*apbE*), molybdenum cofactor (*moeA*, *moeB*, *mobA*), cobalamin (*yvqK*), as well as the Fe-S cluster biosynthesis enzyme cysteine desulfurase (*sufS*). It also includes transporters and secretion system genes (Figure 3H, *btuF*; *crcB*; *mscL*; *mscS*; *gspE*; *comEA*; *comF*).

The enrichment of cofactor and vitamin biosynthesis genes in the Accelerating cluster (Figure 3) prompted us to examine whether cofactor usage itself differs across rate-trajectory clusters. We retrieved representative sequences from *Escherichia coli*, *Bacillus subtilis*, and *Thermus thermophilus* for each LBCA gene and extracted cofactor annotations from UniProt (UniProtConsortium 2025) (Figure 4).

**Figure 4.**
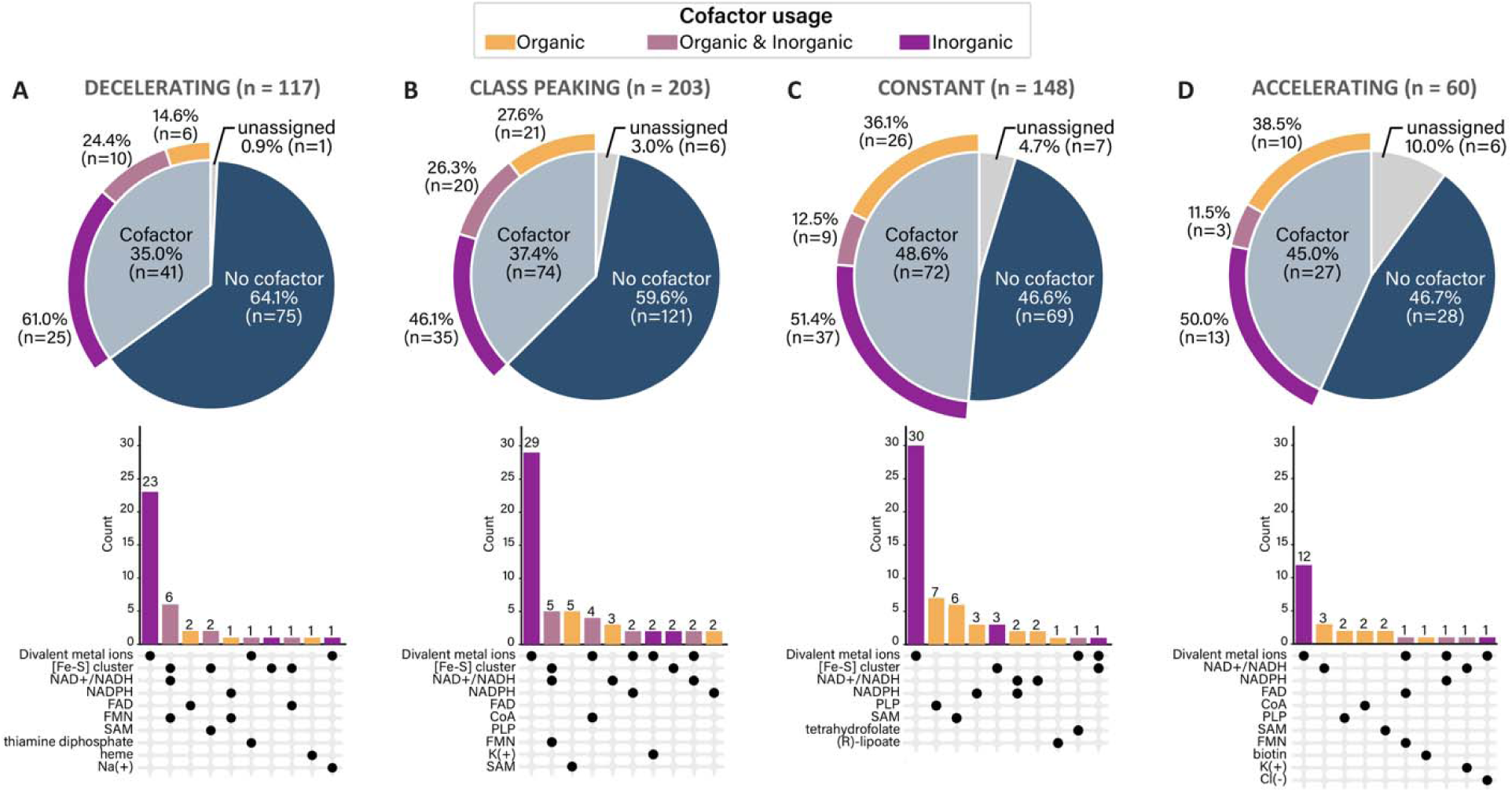
Cofactor usage across evolutionary rate-trajectory clusters. Pie charts show the proportion of cofactor usage, no cofactor requirement, and no cofactor annotation. Cofactor usage is divided into three categories: inorganic only, organic only, or both inorganic and organic. Upset plots below each pie chart show the ten most frequent specific cofactor types and their combinations within each cluster. Clusters are: (A) Decelerating (n=117), (B) Class-Peaking (n=203), (C) Constant (n=148), and (D) Accelerating (n=60).

Among cofactor-using proteins, the proportion relying on inorganic cofactors exclusively or in combination with organic cofactors is highest in the Decelerating cluster (74%), followed by Class Peaking (62%), Constant (61%), and Accelerating (56%). In contrast, the proportion of genes using exclusively organic cofactors increases progressively from Decelerating (26%) through Class Peaking (38%) and Constant (40%) to Accelerating (44%). A higher proportion of exclusively inorganic cofactor usage is most pronounced in the Decelerating cluster, whose genes show the deepest signals of evolutionary divergence.

## Discussion

Evolution of a given gene is typically described by a single rate across evolutionary depth (Wilson, et al. 1977; Jordan, et al. 2002; Rocha and Danchin 2004). This approach is informative but obscures temporal variation of rates. By determining normalized branch lengths across five taxonomic splits (phylum, class, order, family, and genus), our model shows variability of evolutionary rates suggesting changes in selective pressures across time and across the bacterial genome. The 528 genes comprising the LBCA repertoire form a spectrum of rate trajectories. Four clusters of trajectories (Decelerating, Class Peaking, Constant, and Accelerating) capture the major patterns within this continuum. To understand the biological basis of these dynamics, we analyzed gene function and cofactor usage across genes in each cluster.

### The early maturation of the expressome

The Decelerating cluster is enriched in the core machinery of the central dogma, and includes transcription apparatus components, and translation system genes. The pronounced early rate peak observed in the Decelerating cluster highlights an epoch of rapid gene diversification at the phylum and class levels (∼1.5–3.0 Gya (Moody, et al. 2024; Davin, et al. 2025)), followed by a drop in evolutionary rate (Figures 3A-B and 3E). This cluster features all proteins that assemble the expressome, which is the molecular complex that couples transcription and translation in bacteria (Demo, et al. 2017; Kohler, et al. 2017) through physical interactions between transcription factors NusA and NusG, DNA-directed RNA polymerase (RNAP), and several ribosomal proteins (bS1, uS2, bS18, bS21, uS3, uS5, uS10). This pattern extends beyond the expressome to include bacterial-specific translation factors such as IF3, EF-P, and RF3.

Distinctly bacterial innovations in translation underwent an early period of change. During early evolution, genetic information processing genes underwent extensive sequence divergence and biospheric networking via horizontal transfer (Vetsigian, et al. 2006; Bowman, et al. 2026; Kaçar, et al. 2026). Yet, horizontal exchange depends on tree-of-life-wide compatibility, imposing stringent evolutionary constraints to maintain functional interoperability. Once translation initiation and the coupled expressome established in bacteria, substitutions at interaction interfaces between the ribosome and RNAP became overwhelmingly deleterious, effectively locking in sequence identity deep in bacterial history, marking a defining moment in the divergence between bacteria and archaea.

### Core biosynthetic genes diverge at an intermediate evolutionary depth

The Class-Peaking cluster is enriched for enzymes that generate the building blocks of life: nucleotides, amino acids, and membrane lipids. These genes start with lower rates at the phylum level, rise to a peak at the class split, and then decrease markedly. This pattern suggests that strong purifying selection initially constrained these genes but was transiently relaxed around the time of class-level divergences before reasserting at subsequent levels. Overall, the fundamental difference between metabolic supply pathways (concentrated in the Class-Peaking cluster) and the informational systems they feed (concentrated in the Decreasing cluster) is that metabolic enzymes start with low proportional change while informational proteins start high. Although biosynthetic genes are less constrained by physical interaction networks than the ribosome or replisome, they are constrained by the tightly balanced stoichiometry of metabolism.

### Planetary geochemical evolution drives a metallocofactor paradox

Genes utilizing metal cofactors are enriched in the Decelerating and Class-Peaking clusters, whereas metallocofactor biosynthetic pathways (such as heme, molybdopterin, cobalamin, and Fe–S cluster assembly) reside predominantly in the Constant and Accelerating groups. Thus, the rate trajectories of proteins that use inorganic cofactors (Raymond and Segrè 2006; Anbar 2008; Glass, et al. 2014) are decoupled from those of proteins that synthesize organic cofactors and metal clusters (Braakman and Smith 2013). Consistent with this pattern, energy metabolism genes, which depend on iron-sulfur clusters and metal cofactors for function, are concentrated in the Decelerating cluster.

These differences in rate trajectories provide evidence for planetary co-evolution during and after the Great Oxidation Event (Anbar 2008; Lyons, et al. 2024; Robbins, et al. 2026). In the ancient, anoxic oceans preceding and accompanying early bacterial diversification, reduced transition metals (e.g., Fe^2+^, Mn^2+^) were highly soluble and bioavailable. Proteins that directly incorporate these ambient inorganic catalysts adapted as soluble iron and other trace metals precipitated into insoluble oxides when atmospheric and oceanic oxygenation progressed during and after the Great Oxidation Event. This environmental shift exerted strong selective pressure on bacterial enzymatic pathways for cofactor biosynthesis, and for metal mobilization and scavenging (e.g., transporters in the Accelerating cluster). Thus, while the rates of core metalloproteins decelerated early, the biosynthetic pathways required to supply their cofactors maintained higher rates of change as life adapted to post-GOE geochemical environments. This pattern is specific to cofactor biosynthesis and not a general property of biosynthetic pathways, since nucleotide, amino acid, and lipid biosynthetic genes are concentrated in the Class-Peaking cluster.

### Late acceleration and ongoing ecological adaptation

The Accelerating cluster represents genes under continuous rates of change throughout bacterial evolution. The enrichment of transporters, mechanosensitive channels (MscL, MscS), natural competence machinery (ComEA, ComF), and fluoride exporters (CrcB) in this group underscores the role of surface-exposed and environmental-interface proteins in niche adaptation. As bacteria diversify and adapt, genes involved in environmental sensing, osmotic protection, xenobiotic defense, and horizontal gene acquisition remain evolutionarily fluid, accumulating substitutions at higher relative rates near family and genus divergences.

Together, these findings demonstrate the imprints of major molecular constraints and planetary geochemical shifts on the evolutionary history of the LBCA gene repertoire. This implies that the deep conservation typically emphasized in LBCA reconstructions is not uniform, reflecting ongoing adaptation to shifting nutrient availability, environmental redox state, or metabolic niche across the radiations.

## Methods

### Data acquisition

To assess the full breadth of bacterial and archaeal diversity, we downloaded all prokaryotic genomes and taxonomic information from NCBI (Sayers, et al. 2025). Taxonomic information spanning superkingdom to strain level was appended to each genome. We filtered genomes, retaining only those with defined genus designations in RefSeq and passing NCBI’s ANI-based taxonomy check and excluded “Inconclusive” or “Failed” assignments as well as single-cell or metagenomic assemblies. We then selected a single representative genome from each genus, prioritizing complete genomes over scaffolds and contigs, with preference for larger assemblies when completeness was equivalent.

### Gene finding and homologous gene clustering

To ensure uniform data quality, we performed gene finding on all raw assembly files using GeneMarkS-2 (Lomsadze, et al. 2018). We then clustered homologous genes using a divide-and-conquer approach to manage the extensive diversity of bacterial genera (n=2,982). We evenly divided genomes from larger bacterial taxonomic classes (>20 genera, n=18 classes) into ten subsets. Each subset was compared to a control dataset comprising all archaeal genera (n=144) and all bacterial genomes from smaller taxonomic classes (<20 genera, n=88 classes).

We performed all-versus-all sequence similarity searches using DIAMOND v2.1.8.162 (Buchfink, et al. 2021) with an e-value threshold of 10⁻□, option k=0 to retrieve all hits, and the ‘--very-sensitive’ flag to capture distant homologs with <40% identity. We clustered homologous genes using the Markov Cluster (MCL) algorithm (Van Dongen 2008) with a scheme of 2 and inflation parameter of 1.3, which balanced accurate clustering of distantly related sequences against false positives (Brohée and van Helden 2006).

We merged the ten comparisons using a custom Python script that systematically identified clusters assigned to each control dataset gene across the subsets, then merged these clusters. To confirm accuracy and reduce potential over-clustering, we reanalyzed each merged cluster using MCL under the same specifications. This reanalysis either subdivided merged clusters into smaller, more clearly defined groups or confirmed them as singular homologous gene clusters. Each cluster was annotated using eggNOG-mapper v2 (Cantalapiedra, et al. 2021).

To address the imperfect nature of any clustering approach, we performed supplementary clustering using significantly relaxed parameters (scheme 7, inflation parameter 0.2). We manually compared our original dataset to these relaxed clusters and to the 298 genes identified by Moody et al. 2024 (Moody, et al. 2024) from seven previously established universal gene sets, merging clusters where consensus of universality existed across approaches.

### Alignment and phylogenetic analyses

We identified gene clusters as bacterial-specific (present in >75% of bacteria, <10% of archaea) or universal (>75% of both domains). Clusters with average homologous gene copy numbers >2 per genome were excluded. We aligned the remaining clusters using Clustal-Omega with a cluster size of 100 and 10 iterations (Sievers and Higgins 2018).

To determine an appropriate substitution model, we ran ModelFinder analysis within IQ-TREE using the universal ribosomal protein uL16 from both archaea and bacteria combined as a benchmark (Kalyaanamoorthy, et al. 2017; Minh, et al. 2020). We built gene tress using IQ-TREE with 1,000 ultrafast bootstrap replicates under the LG+R10 substitution model (Minh, et al. 2013; Hoang, et al. 2018; Minh, et al. 2020).

Accurate comparison of homologous gene phylogenies required pruning to only single-copy orthologs. We determined the species phylogeny using ASTRAL-Pro3 and all universal and bacterial-specific gene phylogenies (Zhang and Mirarab 2022). For taxa possessing multiple homologous gene copies per phylogeny, we used the Evolutionary Similarity Index (ESI) to select which copy best reflected the species phylogeny. This approach uses pairwise distance matrices between phylogenies to select the gene copy resulting in the minimum sum of squared residuals (Rangel, et al. 2021).

### Evolutionary rate trajectory analyses

To identify each gene’s evolutionary rate trajectory across deep time, we calculated normalized branch lengths at different taxonomic depths for all bacterial-specific and universal genes. Our goal was to quantify the relative evolutionary change occurring in each gene leading up to major taxonomic divergences.

To ensure evolutionary rate patterns reflected genuine biological signal rather than alignment saturation artifacts, we re-estimated branch lengths using only the slowest-evolving alignment sites. For each gene phylogeny built with the LG+R10 substitution model, we extracted posterior mean site rates assigned by IQ-TREE. The LG+R10 model classifies sites into 10 rate categories, ranging from nearly invariant (category 1) to rapidly evolving (category 10). Fast-evolving sites accumulate multiple substitutions at the same position that erase historical signal, while slow-evolving sites maintain phylogenetic information over deep time.

For each gene alignment, we constructed a filtered alignment containing only sites assigned to rate categories 1-5 (the slowest 50% of sites). We used IQ-TREE to re-estimate branch lengths on the original phylogenetic topology (held fixed using the - te option) with the filtered alignment. This approach preserves phylogenetic relationships inferred from the full alignment while calculating branch lengths based exclusively on sites least affected by saturation.

Because gene phylogenies contain multiple representatives per taxonomic group, we used a bootstrap resampling approach for robust median branch length estimates. For phylum, class, order, and family taxonomic levels, each gene tree was pruned to one representative per taxonomic group at the level being measured and immediately above it, isolating the branch spanning that single interval (e.g. phylum-to-class divergences). After pruning, each terminal branch represents the length from the Most Recent Common Ancestor (MRCA) of the higher-level group to the MRCA of the nested lower level group. For example, the phylum-to-class branch runs from a phylum’s MRCA to the MRCA of the classes within it, capturing total evolutionary change accumulated during that interval only (not cumulative from the LBCA root). For each taxonomic level, we took the median length across all sampled groups at that level. To ensure robustness, we repeated this pruning process 100 times for each taxonomic level, each time selecting different representatives. We took the median of all 100 median branch length replicates as the final value for that taxonomic level for each gene.

For each replicate, we assigned one genus per taxonomic group. Candidates were selected based on two criteria: minimal reuse across previous replicates and maximal taxonomic distance from genera already assigned within the current replicate. Taxonomic distance between two genera was calculated as the number of taxonomic levels at which they differ (across phylum, class, order, and family), ranging from 0 (identical taxonomy) to 4 (differing at all levels). Candidates were ranked by usage count (ascending), then by the sum of taxonomic distances to already-assigned genera (descending). The highest-ranked available candidate was selected for each taxonomic group in each replicate, ensuring each replicate sampled broadly across available taxonomic diversity while avoiding overrepresentation of any lineage.

For genus-level branches representing the most recent divergences, we calculated the median terminal branch length directly from the 100 bootstrap replicate phylogenies generated during IQ-TREE analysis with branch lengths adjusted according to the slowest alignment sites, without additional pruning. Domain and kingdom taxonomic groups were excluded due to insufficient sample sizes. Species taxonomic groups were excluded as well because our genome sampling only included one taxon per genus.

### Clustering and principal coordinate analysis

Evolutionary rate trajectories were clustered using k-means clustering (k=4) to identify and characterize distinct temporal patterns. All clustering analyses used scikit- learn’s StandardScaler for feature normalization and KMeans implementation with Euclidean distance metrics (random_state=42, n_init=10) (Pedregosa, et al. 2011).

We calculated average pairwise amino acid identity for each gene alignment to assess overall sequence conservation. This was computed as the mean proportion of identical amino acid positions across all pairwise sequence comparisons within each alignment using custom Python scripts with Biopython (Cock, et al. 2009).

### Cofactor usage

For each LBCA gene, KO assignments were used to retrieve the gene identifier from one of three reference organisms, in priority order: Escherichia coli K-12 MG1655 (eco), Bacillus subtilis 168 (bsu), or Thermus thermophilus HB8 (tth). KEGG gene identifiers were then mapped to UniProt accessions, from which cofactor annotations were extracted.

## Author contributions

H.B.H. conceived the study, designed and implemented the computational pipeline, performed all phylogenetic and evolutionary rate analyses, conducted the clustering and principal coordinate analyses, performed the protein interaction network analyses, interpreted the results, designed data visualizations, and wrote the manuscript. T.G. contributed to literature validation of the gene clustering and homology detection pipeline. A.S.P. contributed to the clustering analyses and edited the manuscript. L.D.W. supervised the project and edited the manuscript. C.A.C. supervised the project, contributed to the study design and interpretation of the results, and edited the manuscript. All authors reviewed and approved the final manuscript.

## Acknowledgements

We thank Gregory P. Fournier for valuable discussions during the early stages of this project. This work was supported by National Aeronautics Space Agency Grant no. 80NSSC24K0344. CAC is funded by the Royal Society Newton International Fellowship [NIF\R1\231037].

## Supplementary materials

**Supplementary Table 1**: Excel spreadsheet including all metadata associated with LBCA genes and taxonomy of genomes used to create figures within the article.

Raw LBCA gene alignments and phylogenies supporting the conclusions of this article are provided at https://doi.org/10.6084/m9.figshare.31511881.

